# Population-wide copy number variation calling using variant call format files from 6,898 individuals

**DOI:** 10.1101/504209

**Authors:** Grace Png, Daniel Suveges, Young-Chan Park, Klaudia Walter, Kousik Kundu, Ioanna Ntalla, Emmanouil Tsafantakis, Maria Karaleftheri, George Dedoussis, Eleftheria Zeggini, Arthur Gilly

## Abstract

**Motivation:** Copy number variants (CNVs) are large deletions or duplications at least 50 to 200 base pairs long. They play an important role in multiple disorders, but accurate calling of CNVs remains challenging. Most current approaches to CNV detection use raw read alignments, which are computationally intensive to process.

**Results:** We use a regression tree-based approach to call CNVs from whole-genome sequencing (WGS, >18x) variant call-sets in 6,898 samples across four European cohorts, and describe a rich large variation landscape comprising 1,320 CNVs. 61.8% of detected events have been previously reported in the Database of Genomic Variants. 23% of high-quality deletions affect entire genes, and we recapitulate known events such as the *GSTM1* and *RHD* gene deletions. We test for association between the detected deletions and 275 protein levels in 1,457 individuals to assess the potential clinical impact of the detected CNVs. We describe the LD structure and copy number variation underlying the association between levels of the CCL3 protein and a complex structural variant (MAF = 0.15, p = 3.6×10^-12^) affecting *CCL3L3*, a paralog of the *CCL3* gene. We also identify a *cis-*association between a low-frequency *NOMO1* deletion and the protein product of this gene (MAF = 0.02, p = 2.2×10^-7^), for which no *cis-* or *trans-* single nucleotide variant-driven protein quantitative trait locus (pQTL) has been documented to date. This work demonstrates that existing population-wide WGS call-sets can be mined for CNVs with minimal computational overhead, delivering insight into a less well-studied, yet potentially impactful class of genetic variant.

**Availability:** The regression tree based approach, UN-CNVc, is available as an R and bash executable on GitHub at https://github.com/agilly/un-cnvc.

**Supplementary Information:** Supplementary information is appended.

## Introduction

Up to 19.2% of the human genome is susceptible to copy number variation, which can have a severe impact on gene function(Zarrei, et al., 2015). Whole-genome sequencing (WGS) at high depth has been the gold standard for detecting large polymorphisms. However, calling structural variants genome-wide has been an ongoing challenge throughout the history of computational genetics. So far, even recent WGS-based structural variant studies are usually made in a limited number of samples or concentrated on targeted regions of the genome(Kayser, et al., 2018; Lu, et al., 2017; Zarrei, et al., 2015). This is because detecting structural variants requires a different study design compared to association studies: whereas for the latter, haplotype diversity and hence sample size are key (Alex Buerkle and Gompert, 2013; Le and Durbin, 2011), for the former, high depth of sequencing is paramount, leading to prohibitive costs for population-wide studies. Structural variant detection also poses a computational challenge, since most algorithms use aligned reads or read pileups as a starting point for event detection. As these file formats describe the entire read pool, processing them genome-wide across an entire population with high-depth WGS is demanding both in terms of running time and memory. In contrast, detecting deletions and insertions from existing variant call sets demand much less compute effort. Such methods were pioneered in the era of genotyping chips (PennCNV (Wang, et al., 2007) and PlatinumCNV (Kumasaka, et al., 2011)), are still widely used (Kayser, et al., 2018; Selvanayagam, et al., 2018) and have recently been proposed to call CNVs from marker-level data in paired cancer samples (Putnam, et al., 2017). To our knowledge, no such method exists for variant calls produced from population-scale whole-genome sequencing (do Nascimento and Guimaraes, 2017). Such variant call sets are typically produced in the Variant Call Format (VCF) in most association-focused studies, and analysis of these comparably small files for CNV calling would be computationally efficient. Here, we evaluate the effect of copy number variants on sequencing depth measured at variant sites using a novel tool (UN-CNVc), and provide a proof-of-concept for calling these large variations in population-wide WGS variant call sets.

## Materials and Methods

The observed read depth for a single sample in a WGS experiment can be modelled as a noisy piecewise constant function: 
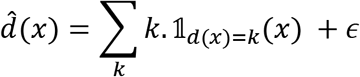
where *d*(*x*) = 0.5*n* is the ideal relative depth at position *x*, *n* is the copy number at this position, 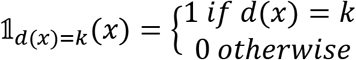 is the indicator function for copy number *k* genome-wide and 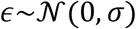 is the error in estimating true read counts. This error term captures all non-CNV factors influencing read depth, such as GC content or reference sequence quality. These variations tend to act on a short range, and over long stretches of sequence, average depths vary little around the per-sample mean (Supplementary Figure 1).

Methods for fitting piecewise constant functions for CNV detection have included circular binary segmentation(Olshen, et al., 2004; Venkatraman and Olshen, 2007), hidden Markov models (Seiser and Innocenti, 2014), smoothing approaches (Hsu, et al., 2005; Tibshirani and Wang, 2008) as well as Bayesian methods(Hutter, 2007), often in the context of array comparative genomic hybridization studies. Here, due to the density of the input dataset, we use regression trees to fit a piecewise constant function, although any segmentation algorithm able to handle hundreds of thousands of points could be used instead. Regression trees have been applied to WGS-based detection of CNVs before(Chen, et al., 2015), and they have been used in analysing variant-level data from paired cancer samples(Putnam, et al., 2017). We wrote the Unimaginatively Named CNV caller (UN-CNVc), a simple and fast CNV detection tool based on regression trees. Due to its sparse input format and the simplicity of the model used, it is able to process call sets from thousands of samples with WGS data in reasonable time. A summary of the CNV calling pipeline is described in Figure 1.a.

**Figure 1:**
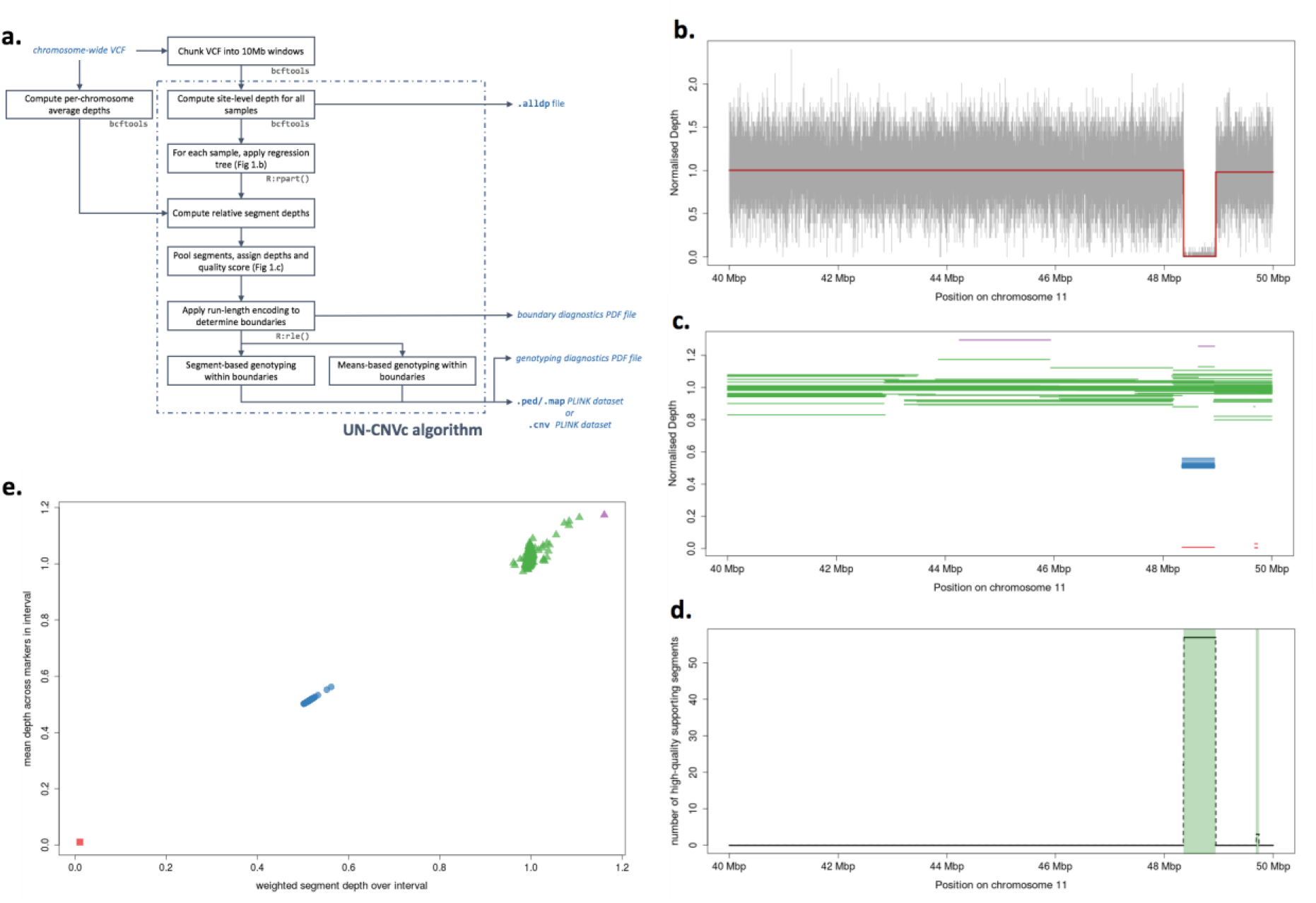
Overview of the UN-CNVc algorithm. **a.** Overview of the pipeline, with input and output files in blue, and external tools and libraries in grey. **b**. Output of a piecewise constant regression (in red) on a 10Mb window on chromosome 11, for a homozygous deletion carrier. The gray signal is the raw relative depth at every sequenced marker for that sample. **c**. Pooled regressed segments across the population, with colour indicating the attributed ideal depth (0:red, 0.5:blue, 1:green, 1.5:purple). **d**. Raw count (dashed line) and run-length encoding (shaded green bars) on the number of high-quality segments with ideal depth < 1. **e**. Genotyping using both weighted average segment depth (colour, scheme identical to **c**.) and average depth across markers (plotting glyphs, squares: 0, circles:0.5, triangles:1). Segment-based genotyping tends to be biased towards the reference for smaller events, whereas means-based genotyping is agnostic to variant size. Both methods tend to perform equally well for large variants.

### Identifying variant regions

Briefly, for each sample in 10Mb windows spanning the entire genome, we apply a regression tree using the rpart R library to the depth at marker sites normalised by chromosome-wide depth. We use the default values of 0.01 for the complexity parameter of the regression tree (the overall *r^2^* of the model must increase of at least this value at each iteration) and 6 for the minimum leaf size. At sample sizes expected in cohort-wide WGS data (>100) in 10Mbp windows, these parameters are very restrictive, i.e. they will only fit a model that follows very broad variations of the data (Fig 1.b). Assembling the constant segments of depth across the entire set of samples provides a global picture of broad depth changes in each 10Mb window (Fig 1.c). Despite an apparent wide diversity of observed depths, the regressed segments cluster around multiples of 0.5 relative depth, as expected if these anomalies indeed corresponded to copy number variants (Supplementary Figure 2).

For each window, we fit a Gaussian mixture model, with means constrained to multiples of 0.5 within the observed depth range at that region. For each depth segment produced by the regression, we assign an ideal depth which is the multiple of 0.5 relative depth that is closest to the actual value of the segment. We also assign a score *s = 2p*, where p is the one-sided p-value for the Gaussian component centered around the ideal depth for that segment, and consider a call high-quality when *s>0.1*. We discretise the window in 5kbp chunks, and consider a chunk as supporting a depth anomaly if the ratio of high-quality versus low-quality segments whose assigned depth is not 1 is greater than 1. To determine boundaries, we then apply run-length encoding (RLE) to this variable, which produces regions in which a majority of high-quality segments support a depth anomaly (Figure 1.d). Application of this method on high-depth WGS data suggests that duplications may exhibit more complex depth variations than deletions. We therefore also implement a deletion-only mode, where only those segments that support deletions are used to call events.

### Segment-based genotyping

Because copy number events can be complex, it is common for a sample to have several segments, and hence several assigned depths per variable region. To produce a single genotype per individual, we compute the mean of the assigned depths weighted by the length of each segment, which is rounded to the next multiple of 0.5. Similarly, we produce an aggregate score summarising the average quality of the regressed segments for that sample. This allows for the easy application of a quality control (QC) step, whereby genotypes with too high a number of segments, or too low an aggregate quality can be set to missing.

### Means-based genotyping

The ability of the regression tree to correctly detect drops or increases in depth depends on the number of markers spanned by a CNV, as well as on the complexity parameter: for a constant complexity, smaller events are harder to distinguish from noise, hence harder to detect. At the limit of detection, it is therefore possible that not every carrier sample exhibits abnormal depth segments, leading to correct calling of the presence of a CNV, but false negative errors in genotyping. To address this issue, we implement means-based genotyping, where each sample gets assigned the multiple of 0.5 that is closest to the average depth across all markers spanning the CNVs called by the regression step (Figure 1.e). The quality score is then simply the distance between the average and assigned depths. This genotyping method is sensitive to incorrect calling of CNV boundaries, but it can perform well on smaller events where segment-based genotyping is inaccurate. We implement a manual genotyper, which applies means-based genotyping on genomic coordinates specified by the user.

## Results

### CNV calling in 6,898 European samples

We apply UN-CNVc on WGS data from 6,898 samples across four studies: the MANOLIS and Pomak isolated cohorts from the HELIC study, the TEENAGE cohort of Greek adolescents, and the INTERVAL study of blood donors in the UK. A total of 401, 353, 349, and 973 CNVs were called from each cohort, respectively. A summary of sample sizes and sequencing depth for each group is given in Supplementary Table 1.

The genome was divided into 332 equal-sized 10 Mbp chunks, which were run in parallel, with some chunks empty due to overlap with pericentromeric regions. Runtime had a power dependency to sample size, between linear and quadratic (Supplementary Figure 3.a) with the linear model giving 2.4 seconds/sample (the best fit was for a *n*^*1.5*^ dependency). On a cluster providing 332 threads, this means UN-CNVc can call CNVs genome-wide on a 1,000-sample cohort in 40 minutes. Peak RAM usage was between a square and a cubic function of the sample number, with approximately 10Gb required for 3,000 samples (Supplementary Figure 3.b).

### Quality control

Quality control (QC) of the variants was carried out based on the plots and statistics files generated by UN-CNVc. Variants called within the centromeres and telomeres were first removed due to the low mapping quality in these regions. Following this, two rounds of QC were performed on the remaining CNVs. First, segment or boundary QC excluded variants based on calling metrics and diagnostics plots, with passing events having no multiple breaks within the call regions and homogenous boundaries (Supplementary Figure 4). Second, genotype QC was performed using the genotype diagnostics plots. For complex events with multiple breakpoints, or small events with incorrect genotypes, boundaries were adjusted using the manual genotyper. (Supplementary Figure 5).

Following this QC procedure, we call 1,320 CNVs across the four cohorts (Table 1). Most of the variants that failed the QC were concentrated within pericentromeric and telomeric regions (Figure 2). Assembly exceptions were particularly rich in CNVs, and although depth patterns were usually complex in these regions, manual genotyping allowed to recover and genotype robust deletion signatures. 57%, 60%, 49%, and 69% of high-quality events in each of the respective cohorts, MANOLIS, Pomak, TEENAGE, and INTERVAL, overlapped with at least one variant in the Database of Genomic Variants (DGV). To make comparisons more meaningful, we applied a 80% reciprocal overlap criterion, which avoids counting as overlapping cases where a large event spans a much smaller one in another cohort. 101 (7.7%) of the high-quality CNVs were shared between two or more cohorts, among which 12 were shared between all four cohorts and 37 between at least three cohorts. The largest overlap was between Pomak and INTERVAL, which shared 54 CNVs, followed by MANOLIS and INTERVAL, with 42 CNVs (Supplementary Figure 6).

**Table 1:** Number of CNVs called in each cohort. Both segment-based and genotype QC were performed for all four cohorts.

|  | **MANOLIS** | **Pomak** | **TEENAGE** | **INTERVAL** |
| --- | --- | --- | --- | --- |
| Total called | 401 | 353 | 349 | 973 |
| High-quality deletions | 212 | 228 | 109 | 771 |
| Badly-mapped | 150 | 84 | 197 | 155 |
| Centromeric/telomeric | 53 | 55 | 47 | 77 |

**Figure 2:**
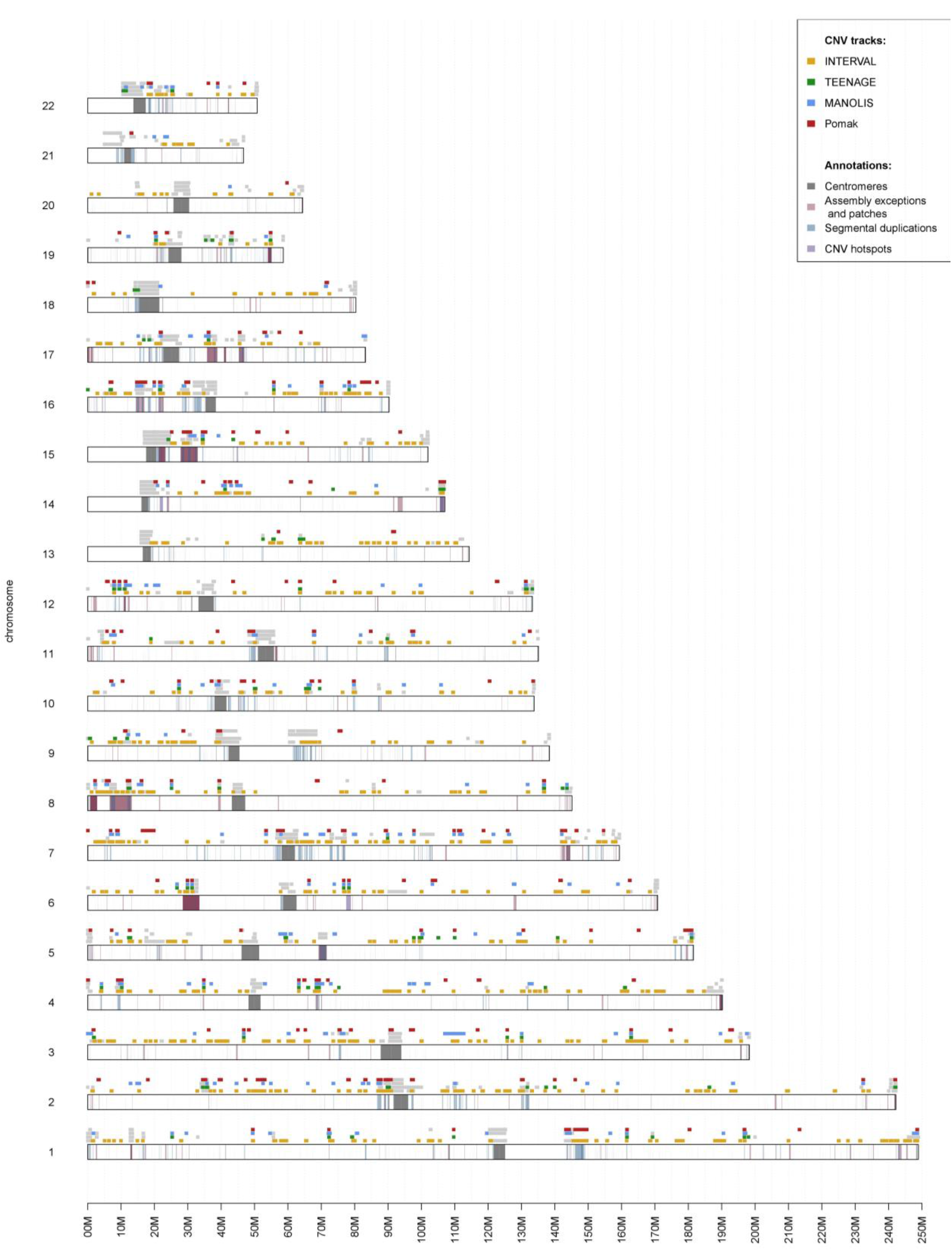
Chromosome map of all CNVs called by UN-CNVc in four cohorts. Light grey tracks represent CNVs that failed QC, while the red, blue, green, and yellow tracks represent high-quality CNVs in MANOLIS, Pomak, TEENAGE, and INTERVAL, respectively. Within the chromosomes, dark grey regions represent the centromeres. Regions marked in pink are assembly exceptions and patches, taken from the GRC data for GRCh38.p12, regions in blue are segmental duplications (from UCSC), and regions in purple are “CNV hotspots”, which are known, highly variable regions comprising an intergenic region on chr6q14.1, an olfactory receptor gene cluster (*OR4C11-OR5L2*) on chr11q11, a leukocyte immunoglobulin gene cluster (*LILRB3-LILRB5*) on chr19q13.42, the immunoglobulin κ, λ, and heavy chain loci (*IGKC, IGLC1, IGH*), and the T cell receptor alpha locus (*TRA*).

We compare the population deletion allele frequencies between any event that was present in at least two cohorts, adjusting for the number of comparisons performed (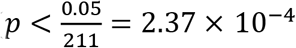 for the two-proportion chi-squared test). We find that 40.5% (41/101) of all shared deletions exhibit significant allelic frequency differences (Supplementary Table 2). The CNV showing the highest heterogeneity in frequency is the known 60kb esv3608493 deletion at 6p22.1, in a region containing 4 *HLA* pseudogenes, *HLA-H, HLA-T, HLA-K*, and *HLA-U*. The deletion occurs most frequently in MANOLIS (MAF = 0.2426) and TEENAGE (MAF = 0.2250), followed by Pomak (MAF = 0.1252) and then INTERVAL (MAF = 0.0987), with the most pronounced difference observed between MANOLIS and INTERVAL (p = 1.68×10^-80^). The low MAF of the variant in INTERVAL corresponds to findings from the Genome Project Phase 3, where frequency in the GBR population was at 0.0879, lower than the European frequency of 0.1113.

### Gene deletions

An average of 51% of our high-quality deletions overlapped protein-coding genes, with 45% of high-quality events deleting at least one exon and 23% deleting one or more entire genes (Supplementary Table 3). Some of these are common deletions that delete genes such as *RHD* and *GSTM1* (Supplementary Text), while a number are in highly-recombinant regions such as the immunoglobulin heavy chain (IGH) locus on chromosome 14q32.33, and are unlikely to be functional. Additionally, we detect a known 58kb deletion overlapping the *BTNL8* and *BTNL3* genes that has been previously predicted to generate a fusion BTNL8/3 protein product (Aigner, et al., 2013) (Supplementary Text). We also find evidence of known disease-associated gene deletions in our cohorts, such as a common 30kb deletion of *APOBEC3B* (chr22:38982347–38992804) that has been associated with increased risk of lung cancer, prostate cancer, (Gansmo, et al., 2018), breast cancer (Han, et al., 2016; Long, et al., 2013; Xuan, et al., 2013) and HIV-1 susceptibility(Singh, et al., 2016), as well as a common CNV at the *FCGR3B* locus (1:161623196–161631963) linked to autoimmune disease susceptibility (Fanciulli, et al., 2007) and malaria severity(Faik, et al., 2017).

### Association analysis

In the MANOLIS cohort, 275 quantitative proteomic traits were assayed using the Proximity Extension Assay provided by Olink Proteomics across three protein panels (Cardiovascular II, Cardiovascular III and Metabolism). We carried out association with the deletions called by UN-CNVc using Plink 1.9. We also applied the linear mixed model implemented in GEMMA, where we accounted for relatedness using an empirical kinship matrix calculated on LD-pruned common SNPs genome-wide. Traits were transformed by applying rank-based inverse normal transformation, and adjusted for 6 covariates: sex, age, age-squared, average levels across all proteins, season of the year, and assay plate. 4 signals pass the genome-wide significance threshold (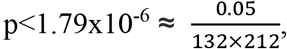 see Supplementary Text). We examined signals down to a suggestive significance level of 1.0×10^-4^ (Supplementary Table 4).

We detect a deletion of the *NOMO1* gene (chr16:14833681–14896160), associated with decreased NOMO1 protein levels (β = –0.6887, σ = 0.1323, p = 2.2×10^-7^). There is no single-point SNV association in that gene for NOMO1 protein levels. The closest SNV association is in the upstream *SHISA9* gene (rs200517050, β = –0.462, σ = 0.0855, p = 1.01×10^-7^), and a stronger association is also present in the *NOMO3* gene (rs3891245, intronic, β = –0.371, σ = 0.0476, p = 5.12×10^-14^). *NOMO1, NOMO2* and *NOMO3* are closely located genes with very high sequence similarity (99.4% and 99.5% homology (BLAST)), and cannot be distinguished by the polyclonal antibody used in the OLINK proteomics assay. Both associations are independent, both of each other (r2<1×10^-3^) and the deletion (r^2^ _rs200517050_ = 0.06, r^2^ _rs3891245_ = 1.2×10^-3^), suggesting that a *NOMO1* deletion and an intronic variant in *NOMO3* independently affect circulating levels of the NOMO proteins.

We find evidence of a complex CNV overlapping the *CCL3L3* gene and influencing CCL3 protein levels (Supplementary Figure 7). We manually genotype a CNV (chr17:36195241–36196130) affecting the last two exons of *CCL3L3*, which is associated with decreased CCL3 levels (MAF = 0.15, β = –0.378, σ = 0.05348, p = 2.55×10^-12^). Copy-number variation of *CCL3L3* and *CCL3L1*, its alias on an alternate haplotype (NT_187661.1) of chromosome 17, have been extensively studied. In addition to levels of their protein product (Townson, et al., 2002), they have been shown to be associated with rheumatoid arthritis(Ben Kilani, et al., 2016; Nordang, et al., 2012), immune reconstitution following HIV therapy (Aklillu, et al., 2013), and protection against malaria (Carpenter, et al., 2012). The gene product of *CCL3L3* binds to the same chemokine receptors as its close paralog *CCL3*, albeit with increased affinity, which suggests that the OLINK proteomics assay might not be able to differentiate between the two ligands. This is even more likely as the two proteins are highly similar in sequence (95% homology; BLAST) and there is no commercially available antibody that can distinguish the two(Carpenter, et al., 2012). Up to 14 copies of *CCL3L3* have been validated in some genomes(Sudmant, et al., 2010), with the majority of people carrying 1 to 6 copies (Rimoin, et al., 2013), whereas we confirm up to 7 copies in the MANOLIS cohort. It has been hypothesised that increased copy number of this gene resulted in higher levels of expression of its protein product, however in our study, including copy numbers greater than 2 in the model weakened the association compared to a deletion-only model (Supplementary Figure 8) suggesting that although deletion of *CCL3L3* decreases CCL3 levels, those levels are not affected by gene duplication.

## Discussion

### Comparison with other callers

We compare UN-CNVc’s calling performance genome-wide with PennCNV, an array-based method, and the CNV discovery pipeline of GenomeStrip, a sequencing read-based method, on 211 MANOLIS samples with both sequencing and CoreExome array data. On this subset, PennCNV took 2 hours to run with 586Mb peak RAM use, and GenomeStrip tool 14.5 hours with peak RAM use of 3Gb, compared to 16 minutes and 798Mb for UN-CNVc. On these samples, UN-CNVc calls 253 CNVs in deletion-only mode, whereas PennCNV and GenomeStrip call 1,404 and 10,660 CNVs with minimum copy number <2, respectively. As expected, our method called on average larger CNVs than the other two methods (Supplementary Figure 9). Only 18 (7%) of UN-CNVc’s events overlapped GenomeStrip’s with 80% reciprocal overlap, however, 138 (55%) further regions called as variable by UN-CNVc completely contained one or more GenomeStrip CNVs. Only 24 (9%) of UN-CNVc’s CNV regions had 80% reciprocal overlap with the CNVs called by PennCNV, while 72 (28%) regions completely contained one or more PennCNV variants.

### Overlap with a database of known CNVs

Compared to the other two methods, a higher percentage of CNVs detected by UN-CNVc had 80% reciprocal overlap with known CNVs in the Database of Genomic Variants (DGV; build 38, May 2016), although PennCNV and GenomeStrip each called more known CNVs. For UN-CNVc, 110 (55%) CNVs overlapped with known variants in DGV, versus 437 (16%) and 2,615 (25%) for PennCNV and GenomeStrip, respectively, indicating that UN-CNVc offers higher specificity and lower sensitivity than the other methods. A similar trend was observed for the number of gene-deleting regions, where 66 (33%) of UN-CNVc deletions affected entire genes, compared to 102 (4%) for PennCNV, and 74 (0.7%) for GenomeStrip. Notably, the complete deletion of the *RHD* gene was detected only by UN-CNVc in the 211 MANOLIS samples. For the array-based PennCNV, this was likely due to the lack of tagging SNPs within the region. Only 5 tagging SNPs in the CoreExome array were within the *RHD* gene coordinates, compared to 141 SNPs from the WGS data used by UN-CNVc, demonstrating the advantage of using WGS data for CNV calling. For GenomeStrip, the deletion was split into six smaller CNVs with an average size of 11kb. This example, where the whole gene is known to be deleted, indicates that in some cases GenomeStrip may be tiling large CNVs by dividing them in smaller events.

### Limits of the piecewise constant regression model

Despite providing a certain level of automation, UN-CNVc still requires post-run manual QC, in the same way as array-based genotypes require inspection of cluster plots. The software generates extensive diagnostic tables and plots to make this task easier for the user. Although piecewise constant regression can accurately model WGS depth in a single individual, UN-CNVc leverages large sample sizes (n>100) to differentiate signal from noise. Furthermore, since the software performs clustering on depth averages, a high enough depth (>15x) is required to ensure proper cluster separation. Finally, using marker-level depth puts limits on the precision of the boundaries as well as the sizes of detected CNVs. The maximum precision achievable by a method such as UN-CNVc is the distance between two consecutive SNVs, in practice it is limited to around 10kb by the minimum leaf size, the segment aggregation algorithm and the discretization step. Our method relies on at least one correct call by piecewise constant regression to genotype a CNV, which makes small, rare CNVs difficult to call.

For cases where the study design deviates from the ideal use case above, users can adjust the sensitivity of UN-CNVc using several parameters. First, the complexity value passed directly to the regression tree directly influences the elasticity of the regression tree model (Supplementary Figure 10). Smaller values allow the piecewise constant regression to follow depth more closely, therefore allowing to detect smaller CNVs but increasing the risk of false positives. This parameter can be adjusted by starting at the default value of 0.01 and decreasing it until a reference deletion (e.g. the *RHD* gene deletion) is correctly detected and the number of carriers stops increasing. Second, the window size, which should be increased from its default of 10Mb if sample size is low (<100). Third, the ratio of high-quality vs. low-quality segments required to call a deletion, which can be increased from its default value of 1 when analysing a particularly noisy depth signal. Fourth, the discretisation step, which is set by default at 5kbp, and which determines the precision of the CNV boundaries. This value should not be smaller than the minimum distance separating two SNPs, and should be kept reasonably large as decreasing it increases execution time linearly. In practice, changing parameters other than the complexity value should not be necessary under most use cases.

## Conclusion

We demonstrate that it is possible to call large CNVs from variant-level WGS depth information in large cohorts. Compared to other methods, UN-CNVc performs well and offers better specificity, although it is limited to large events. As a proof-of-concept, UN-CNVc successfully detects well-known deletions, such as the complete deletions of *RHD, GSTM1* and *CCL3L1*, in 6,898 samples with deep WGS data. We conduct an association study with 272 quantitative protein levels in a set of 1,457 individuals and find two association signals, in which deletion of the *cis* gene caused a significant decrease in the resulting protein levels. These results provide proof of principle for cohort-wide variant-level depth approaches as a platform for discovering disease-associated CNVs and genes. Accurate read-based methods that integrate within standard single-nucleotide variant calling pipelines, such as the one implemented in GATK4, are under active development. UN-CNVc provides a computationally inexpensive means for CNV calling using only the ubiquitously available depth field from Variant Call Format (VCF) files. This approach is much less intensive than read-based re-analysis, and allows quick screening for areas harbouring copy number variation. These regions can then be taken forward for read-level analysis, which will provide base-pair resolution for breakpoints in CNVs of interest.

## Acknowledgments

The authors thank the residents of the Mylopotamos villages for taking part. The MANOLIS study is dedicated to the memory of Manolis Giannakakis, 1978–2010. We would like to thank the Human Genetics Informatics (HGI) group at the Wellcome Sanger Institute, for performing variant calling on the datasets used in this study. This work was funded by the Wellcome Trust [098051] and the European Research Council [ERC-2011-StG 280559-SEPI]. INTERVAL study: Participants in the INTERVAL randomised controlled trial were recruited with the active collaboration of NHS Blood and Transplant England (www.nhsbt.nhs.uk), which has supported field work and other elements of the trial. DNA extraction and genotyping was funded by the National Institute of Health Research (NIHR), the NIHR BioResource (http://bioresource.nihr.ac.uk/) and the NIHR Cambridge Biomedical Research Centre (www.cambridge-brc.org.uk). The academic coordinating centre for INTERVAL was supported by core funding from: NIHR Blood and Transplant Research Unit in Donor Health and Genomics, UK Medical Research Council (G0800270), British Heart Foundation (SP/09/002), and NIHR Research Cambridge Biomedical Research Centre. A complete list of the investigators and contributors to the INTERVAL trial is provided in (Di Angelantonio, et al., 2017). This report is independent research by the National Institute for Health Research. The views expressed in this publication are those of the author(s) and not necessarily those of the NHS, the National Institute for Health Research or the Department of Health. This work was undertaken by Cambridge who received funding from the NHSBT; the views expressed in this publication are those of the authors and not necessarily those of the NHSBT. The TEENAGE study has been supported by the Wellcome Trust (098051), European Union (European Social Fund—ESF) and Greek national funds through the Operational Program “Education and Lifelong Learning” of the National Strategic Reference Framework (NSRF)— Research Funding Program: Heracleitus II, Investing in knowledge society through the European Social Fund. The GATK3 program was made available through the generosity of the Medical and Population Genetics program at the Broad Institute, Inc. The authors thank Kerstin Howe for her insights on assembly exceptions.

## Author Contributions

Software design: AG, DS

Software development: AG

Analysis: GP, AG

Single-point analysis: YCP

INTERVAL data: KW, KK

Greek cohorts sample collection: IN, ET, MK, GD, EZ

Article writing: AG, GP, EZ

Project management: EZ, AG

