## Supplementary Material for "Population-wide copy number variation calling using variant call format files from 6,898 individuals" (from another version of this preprint: version 1, 2018-12-21)

### Supplementary Data

#### Supplementary Text

*Complete gene deletions of* RHD *and* GSTM1

##### I. RHD

UN-CNVc detected a 70kb deletion spanning *RHD* (chr1:25272393-25330445) in the MANOLIS cohort, and we manually genotyped this known variant in other cohorts. The *RHD* gene codes for the rhesus D antigen (RhD), which determines an individual's rhesus blood group. Rhesus-positive (Rh+) individuals express the antigen on the surface of their red blood cells, whereas rhesus-negative (Rh-) individuals do not. In Europeans, deletion of *RHD* is the most common cause of the Rh- phenotype(Flegel, 2007). We detected 122 (8%), 223 (14%), 15 (15%), and 888 (24%) homozygous individuals in MANOLIS, Pomak, TEENAGE, and INTERVAL, respectively. Only self-reported blood groups were available for some individuals in MANOLIS and Pomak, and as a result there was limited agreement between genotype and phenotype, where some individuals were Rh- despite having at least one copy of *RHD*.

| **MANOLIS** | Rh+ | Rh- |
|---|---|---|
| Non-hom | 62 | 5 |
| Hom | 1 | 5 |

| **Pomak** | Rh+ | Rh- |
|---|---|---|
| Non-hom | 70 | 12 |
| Hom | 1 | 10 |

##### II. GSTM1

Complete deletion of *GSTM1* (1:109682397-109702658) is a very common event, with an average allele frequency of 0.73 in Europeans (1000 Genomes Project Phase 3). The CNV was first detected by UN-CNVc in Pomak, following which we manually genotype the deletion in other cohorts. We find that for all of the cohorts, allele frequency was similar to the European average, at 0.73, 0.77, 0.72, and 0.73 for MANOLIS, Pomak, TEENAGE, and INTERVAL, respectively. GSTM1 null frequency was 53%, 61%, 55%, and 58% for the respective cohorts, which is within the 16%-60% range observed in past studies(Garte, et al., 2001; Kasthurinaidu, et al., 2015). The GSTM1 null genotype has been widely studied and associated with increased risk to a multitude of disorders, including, but not limited to hypertension (Petrovic and Peterlin, 2014), atherosclerosis (Grubisa, et al., 2018), Alzheimer's disease (Wang, et al., 2016), and type 2 diabetes(Nath, et al., 2017).

##### I. BTNL8, BTNL3

The formation of chimeric genes is a common CNV consequence. We detect evidence for this in a 58kb deletion (chr5:180948042-181006042) that is shared among all four cohorts and deletes exons of two adjacent genes, *BTNL8* and *BTNL3*. The same deletion (chr5:180948027-181003596) was confirmed in a previous study to generate a fusion BTNL8/3 protein product which affected the expression of genes involved in immune response.

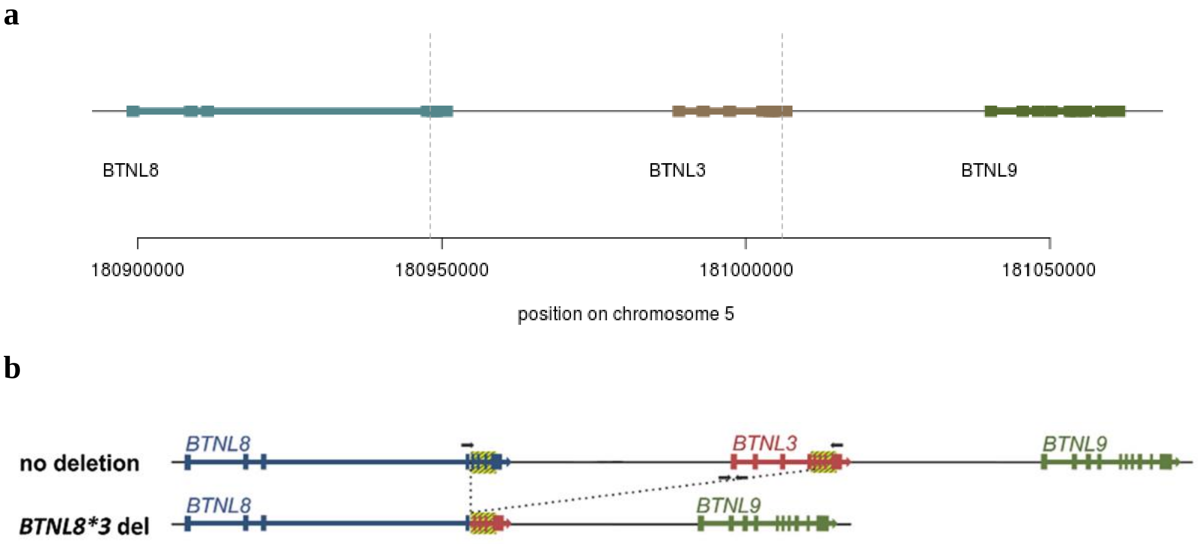


**Deletion spanning exons of *BTNL8* and *BTNL3* results in a *BTNL8/3* chimeric gene. (a)** Grey dotted lines denote the boundaries of the deletion called by UN-CNVc in all four cohorts. The deletion affects the last 4 exons of *BTNL8* and the first 7 exons of *BTNL3*. **(b)** Predicted formation of the *BTNL8/3* chimeric gene according to Aigner *et al.*, 2013(Aigner, et al., 2013). [Figure adapted from Aigner *et al.*, 2013]

*Study-wide significance threshold for association analysis with 275 quantitative proteomics traits*

We establish the study-wide significance threshold p $<1.79\mathrm{x}10^{-6} \approx 0.05$ / (132x212), where 132 is the effective number (Meff) of protein traits and 212 is the number of high-quality deletions in MANOLIS. Meff was calculated using Chevurud's (2001) method(Cheverud, 2001).

#### Supplementary Tables

**Supplementary Table 1: Sequencing characteristics for the four datasets analysed in this study.** For sequencing depth, standard deviations are given in parentheses.

| | MANOLIS | Pomak | TEENAGE | INTERVAL |
|---|---|---|---|---|
| **n** | 1,457 | 1,617 | 100 | 3,724 |
| **Mean depth (SD)** | 22.5 (3.53) | 18.4 (3.54) | 32.2 (2.61) | 18.4 (2.03) |

**Supplementary Table 2:** Significant heterogeneities in frequencies of shared CNVs between cohorts. The significance threshold is $p<2.16\times10^{-4} \approx 0.05/232$, where 232 (10x6 + 24x3 + 50x2) is the number of pairwise comparisons performed.

Link to Supplementary Table 2

**Supplementary Table 3.** Gene-deleting regions in the four cohorts.

| | MANOLIS | Pomak | TEENAGE | INTERVAL |
|---|---|---|---|---|
| **High-quality deletions** | 212 | 228 | 109 | 771 |
| **Gene-overlapping regions** | 116 | 121 | 43 | 397 |
| **Gene-deleting regions** | 52 | 60 | 22 | 172 |
| **Exon-deleting regions** | 105 | 106 | 38 | 349 |

**Supplementary Table 4. Suggestively significant associations in the MANOLIS cohort.** Study-wide significant associations are highlighted in green. Het and hom are the numbers of heterozygote and homozygote carriers in MANOLIS. BETA, SE and P are the effect size, standard error and p-value. For GEMMA, we report the likelihood ratio test p-value.

| | | | | | | Plink 1.9 | | | GEMMA 0.94 | | |
|---|---|---|---|---|---|---|---|---|---|---|---|
| Protein | CHR | START | END | het | hom | BETA | SE | P | BETA | SE | P |
| CCL3 | 17 | 36195241 | 36196130 | 364 | 39 | -0.3781 | 0.05348 | 2.56E-12 | -3.57E-01 | 5.49E-02 | 1.12E-10 |
| TFF3 | 6 | 29881000 | 29939000 | 549 | 79 | 0.2534 | 0.04561 | 3.35E-08 | 2.50E-01 | 4.69E-02 | 1.47E-07 |
| NOMO1 | 16 | 14833681 | 14896160 | 62 | 1 | -0.6887 | 0.1323 | 2.24E-07 | -7.15E-01 | 1.39E-01 | 3.31E-07 |
| TNFR1 | 6 | 31251000 | 31262000 | 155 | 4 | -0.4003 | 0.08286 | 1.51E-06 | -3.48E-01 | 9.05E-02 | 1.33E-04 |
| PPP1R2 | 7 | 53390000 | 53525000 | 74 | 3 | -0.5636 | 0.1182 | 2.06E-06 | -4.68E-01 | 1.23E-01 | 1.64E-04 |
| TYMP | 7 | 53390000 | 53525000 | 74 | 3 | -0.5382 | 0.113 | 2.10E-06 | -4.80E-01 | 1.19E-01 | 6.54E-05 |
| CD1C | 6 | 31269000 | 31271000 | 594 | 760 | 0.1984 | 0.04437 | 8.43E-06 | 1.91E-01 | 4.70E-02 | 5.62E-05 |
| CA13 | 5 | 180948000 | 181005000 | 583 | 125 | 0.1893 | 0.04279 | 1.06E-05 | 1.70E-01 | 4.39E-02 | 1.18E-04 |
| NADK | 7 | 53390000 | 53525000 | 74 | 3 | -0.5019 | 0.1139 | 1.14E-05 | -4.25E-01 | 1.20E-01 | 4.48E-04 |
| PSPD | 10 | 797150000 | 79747000 | 681 | 249 | -0.1717 | 0.03905 | 1.19E-05 | -1.91E-01 | 4.10E-02 | 3.24E-06 |
| CCL24 | 7 | 40205016 | 40295016 | 4 | 0 | 2.514 | 0.5748 | 1.31E-05 | 2.25E+00 | 6.97E-01 | 1.35E-03 |
| NEMO | 7 | 53390000 | 53525000 | 74 | 3 | -0.4965 | 0.1154 | 1.80E-05 | -4.35E-01 | 1.21E-01 | 3.54E-04 |
| MB | 8 | 16080068 | 16150000 | 18 | 0 | -1.148 | 0.268 | 1.96E-05 | -1.12 | 2.66E-01 | 2.93E-05 |
| UPAR | 7 | 53390000 | 53525000 | 74 | 3 | -0.4818 | 0.1132 | 2.24E-05 | -3.86E-01 | 1.20E-01 | 1.42E-03 |
| CDHR5 | 16 | 76920071 | 77015071 | 8 | 0 | 1.581 | 0.3753 | 2.69E-05 | 1.12E+00 | 4.01E-01 | 5.26E-03 |
| CTSL1 | 2 | 84110191 | 84185191 | 12 | 0 | -1.203 | 0.2881 | 3.19E-05 | -1.15 | 3.10E-01 | 2.47E-04 |
| UPAR | 6 | 31251000 | 31262000 | 155 | 4 | -0.3444 | 0.08315 | 3.67E-05 | -3.11E-01 | 9.04E-02 | 6.11E-04 |
| TM | 6 | 29881000 | 29939000 | 549 | 79 | 0.1912 | 0.0462 | 3.71E-05 | 1.90E-01 | 4.93E-02 | 1.23E-04 |
| PAG1 | 7 | 53390000 | 53525000 | 74 | 3 | -0.4732 | 0.1143 | 3.72E-05 | -4.23E-01 | 1.20E-01 | 4.67E-04 |
| CD164 | 5 | 180948000 | 181005000 | 583 | 125 | 0.1749 | 0.04235 | 3.86E-05 | 1.38E-01 | 4.39E-02 | 1.75E-03 |
| ST2 | 22 | 25245222 | 25560222 | 16 | 0 | -1.145 | 0.2778 | 3.98E-05 | -9.89E-01 | 2.87E-01 | 6.18E-04 |
| GRAP2 | 7 | 53390000 | 53525000 | 74 | 3 | -0.6167 | 0.15 | 4.29E-05 | -5.36E-01 | 1.52E-01 | 4.72E-04 |
| SERPINB6 | 6 | 31251000 | 31262000 | 155 | 4 | -0.3349 | 0.08286 | 5.62E-05 | -2.93E-01 | 8.97E-02 | 1.13E-03 |
| CCL15 | 12 | 17500025 | 17650025 | 4 | 0 | 2.024 | 0.5007 | 5.62E-05 | 1.78E+00 | 5.31E-01 | 8.80E-04 |
| IL27 | 8 | 2190000 | 2270000 | 90 | 1 | -0.4441 | 0.11 | 5.74E-05 | -4.32E-01 | 1.20E-01 | 3.54E-04 |
| PILRB | 6 | 31251000 | 31262000 | 155 | 4 | -0.334 | 0.08291 | 5.95E-05 | -3.08E-01 | 9.21E-02 | 8.61E-04 |
| CCDC80 | 7 | 53390000 | 53525000 | 74 | 3 | -0.4539 | 0.1139 | 7.08E-05 | -3.48E-01 | 1.20E-01 | 4.29E-03 |
| FCRL1 | 6 | 103290023 | 103315500 | 724 | 410 | 0.1546 | 0.03883 | 7.26E-05 | 1.10E-01 | 4.05E-02 | 7.15E-03 |
| IL18 | 6 | 73885313 | 73892900 | 578 | 122 | -0.1688 | 0.04278 | 8.37E-05 | -1.75E-01 | 4.43E-02 | 8.24E-05 |
| PRTN3 | 7 | 8790064 | 8830064 | 127 | 4 | 0.3607 | 0.09182 | 9.02E-05 | 2.75E-01 | 9.80E-02 | 5.28E-03 |
| ITGB2 | 7 | 8790064 | 8830064 | 127 | 4 | 0.3605 | 0.09184 | 9.10E-05 | 3.11E-01 | 9.64E-02 | 1.46E-03 |
| MMP9 | 6 | 31251000 | 31262000 | 155 | 4 | -0.3235 | 0.08282 | 9.84E-05 | -2.88E-01 | 9.13E-02 | 1.70E-03 |

### Supplementary Figures

**Supplementary Figure 1: Local variations in depth averaged at variant sites in different window sizes.** Window sizes are in base pairs. Standard deviations are expressed in units of coverage, and are computed on chromosome 11 for one MANOLIS individual. The rightmost point corresponds to the recommended window size of UN-CNVc.

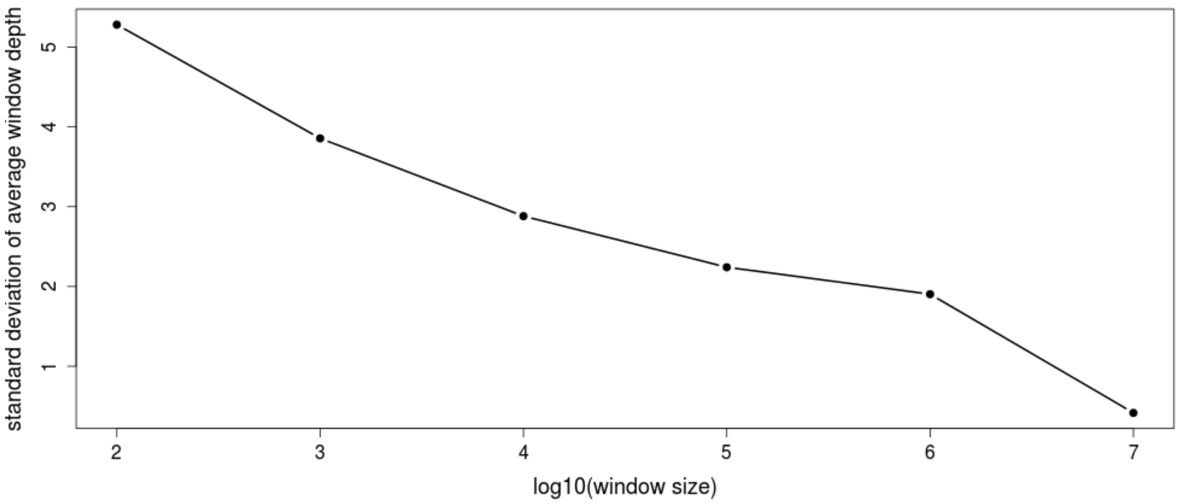

**Supplementary Figure 2: Depth segments after piecewise constant regression prior to QC.** Segment depths are counted for all CNVs called in the HELIC MANOLIS cohort in deletion-only mode. Segments with a called depth of 1 are masked due to their number, so as not to eclipse other depths. Note the clear peaks around 0 and 0.5 for depth segments. Duplication signatures clustering around expected multiples of 0.5 is less apparent, as expected in deletion-only mode.

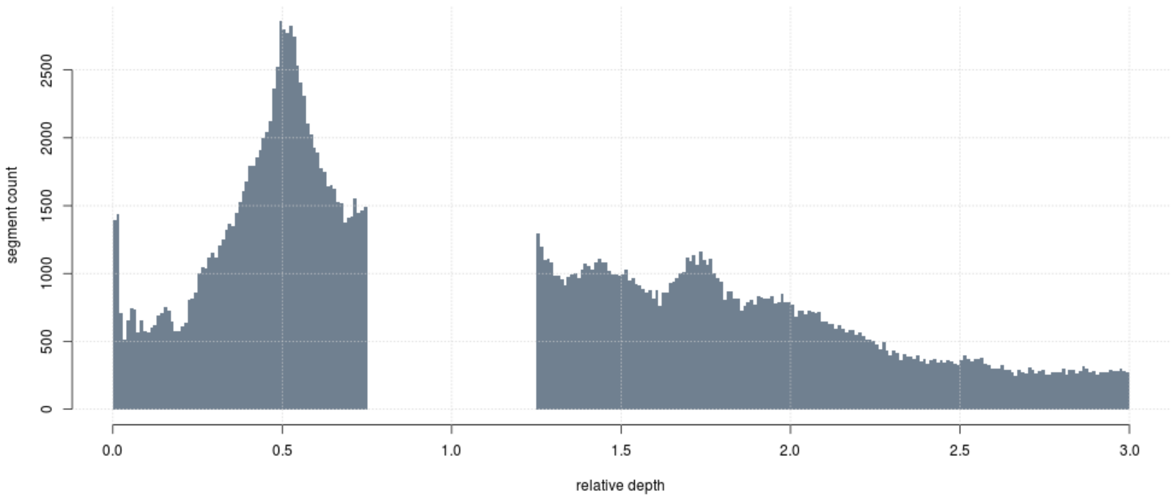

**Supplementary Figure 3: a.** Median runtime as a function of sample size for all three cohorts analysed as part of this study. The black line is a linear model fitted to the data with the intercept constrained to 0. Dashed lines represent the ten $\square = \square^{1+0.1\square}$(n natural>1) power functions between the linear and quadratic models. Red lines represent intervals of one median absolute deviation (MAD) around the median. **B.** Peak RAM usage. Dashed lines represent the four first $\square = \square^{1+0.5\square}$functions (n natural>1).

**a.**

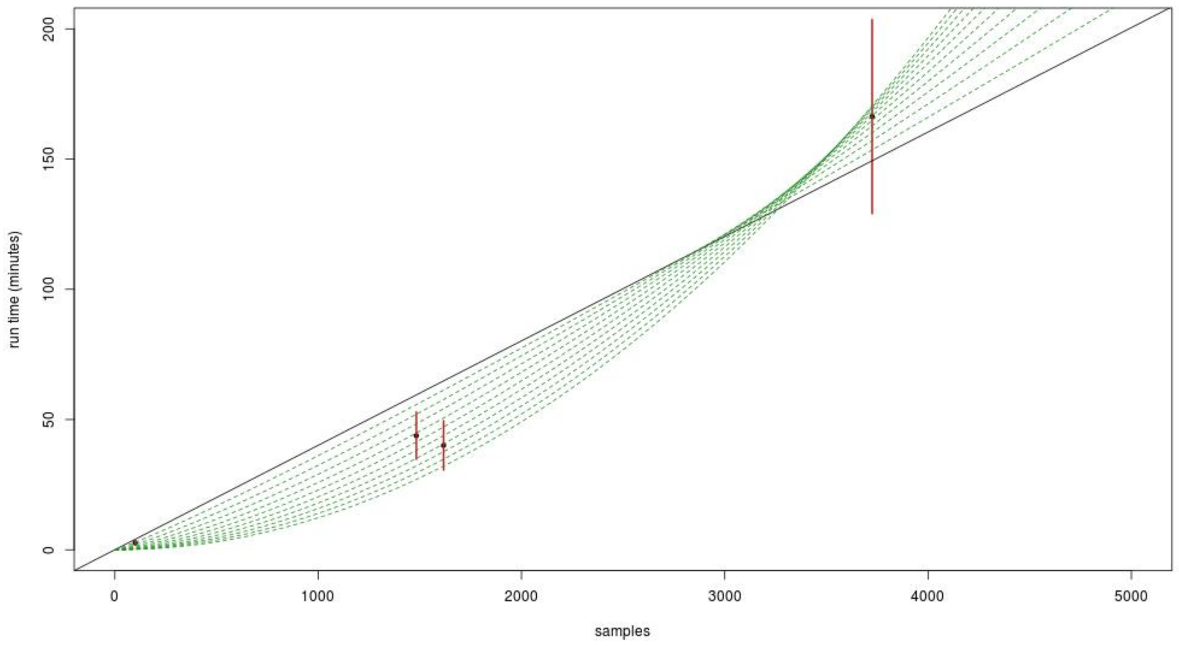


**b.**

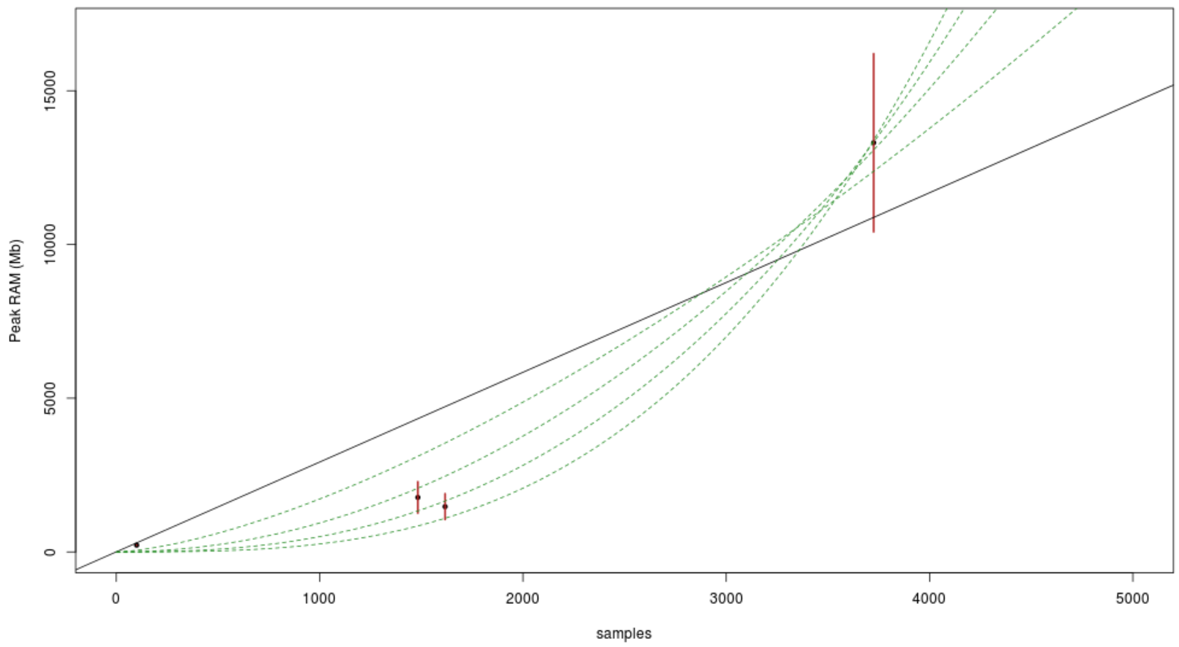

**Supplementary Figure 4: a.** Diagnostic plot of a high-quality and **b.** low-quality deletion based on boundary QC. The top left panel shows the called depth segments in the region with the regions called by RLE in purple. The right panel shows the histogram of weighted segment depths used to fit the Gaussian model. The bottom panel shows the count and RLE graph over the region.

**a.**

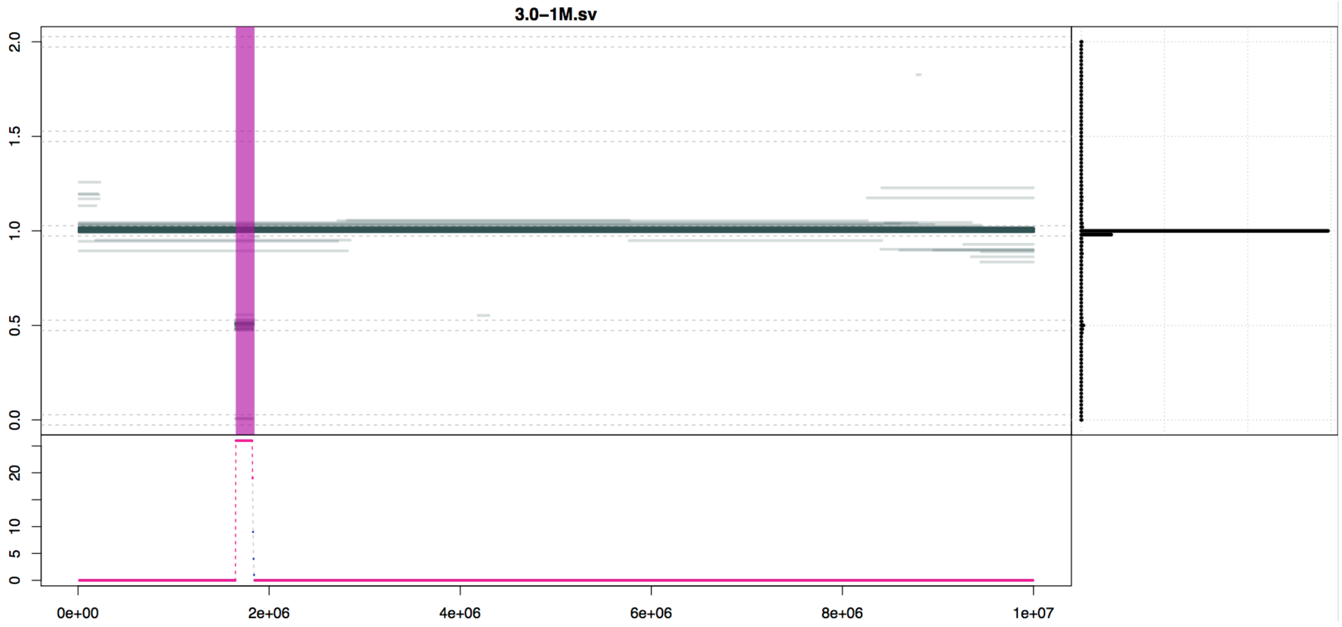


**b.**

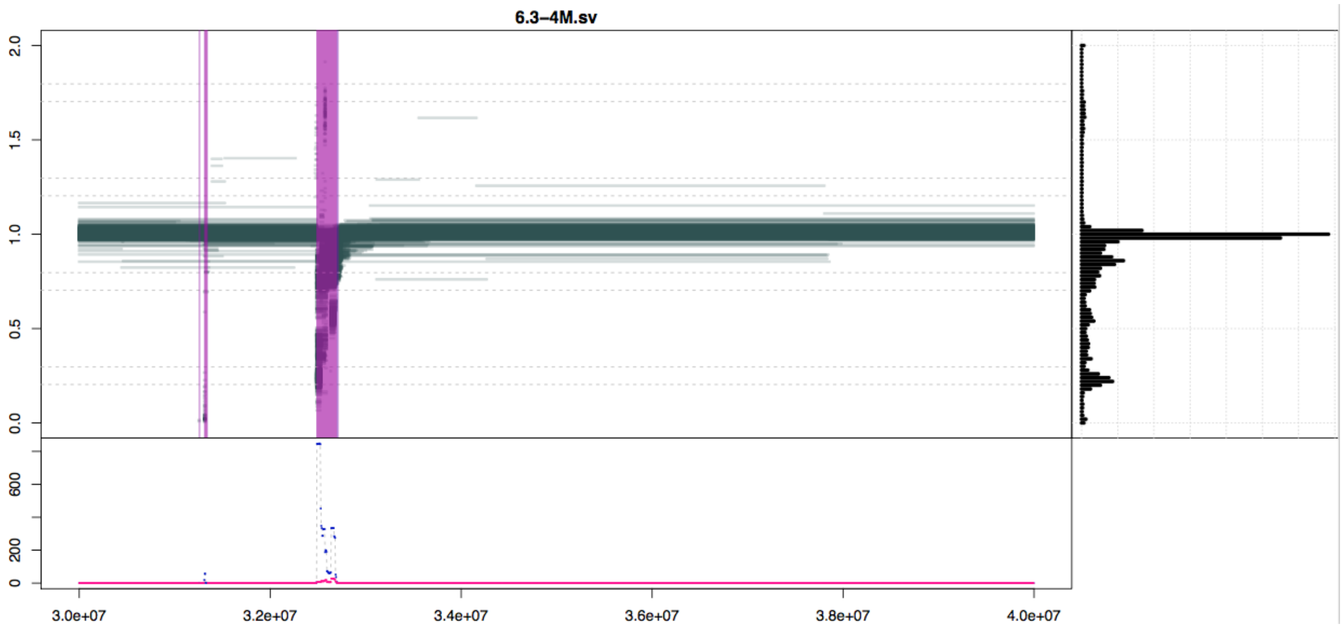

**Supplementary Figure 5: a.** Diagnostic plots of a CNV that requires manual genotyping, based on genotype QC. Top left: average marker depth over called region. Top right: rolling average mean (*k=l/10* where *l* is the length of the called region) for every sample in the region. Bottom right: segments called in the region, with deletion-carrying samples in red. Bottom left: segment (y axis, colour indicates called genotype) and means (x axis, symbol indicates genotype)-based genotype in the region. Means-based genotyping performs better, but overestimates depth for homozygotes. In this example, different segment boundaries have been called in different samples, leading to an overestimation of the deletion size. This has the consequence of biasing depth estimates upwards (top left) with homozygotes incorrectly called as heterozygotes. **b.** Diagnostic plots after manual genotyping. The boundaries and genotypes have been corrected.

**a.**

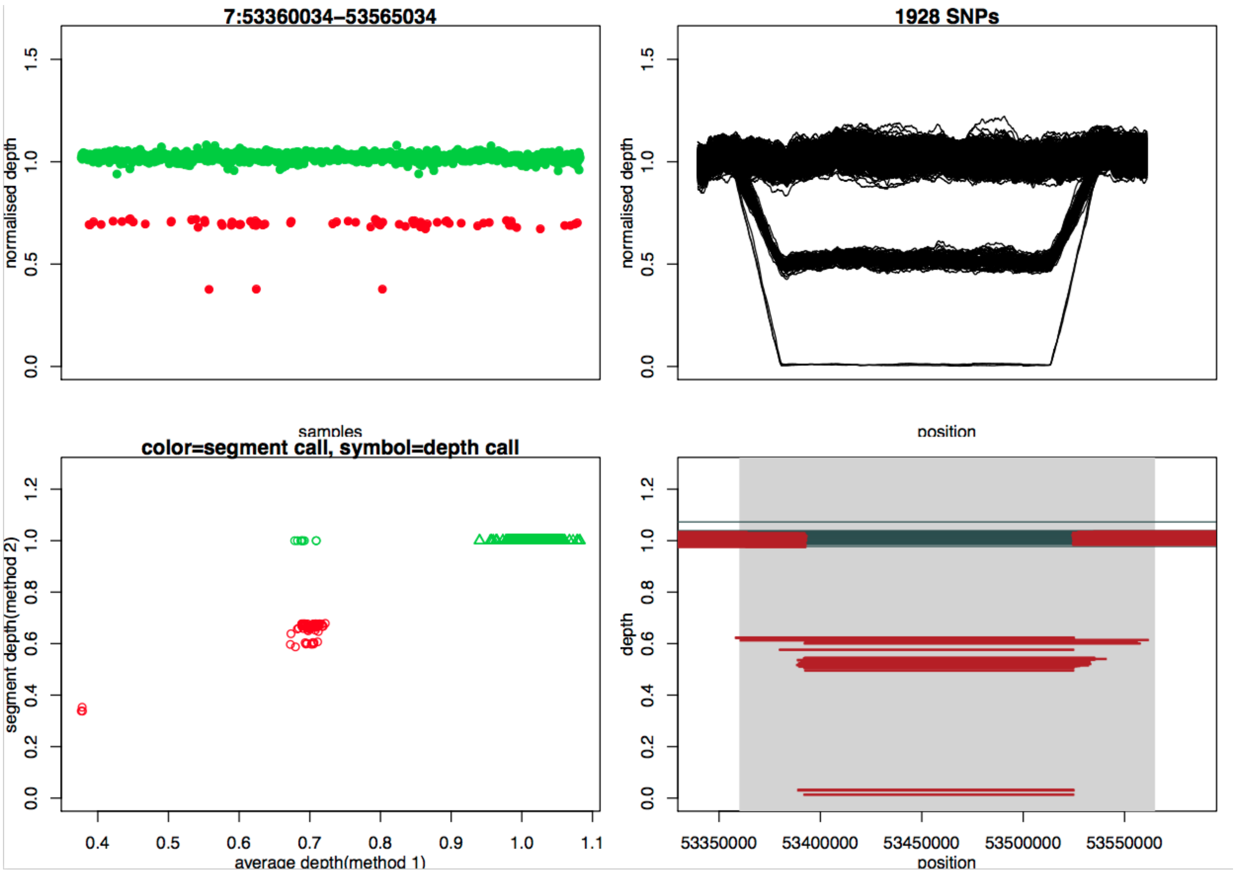

**b.**

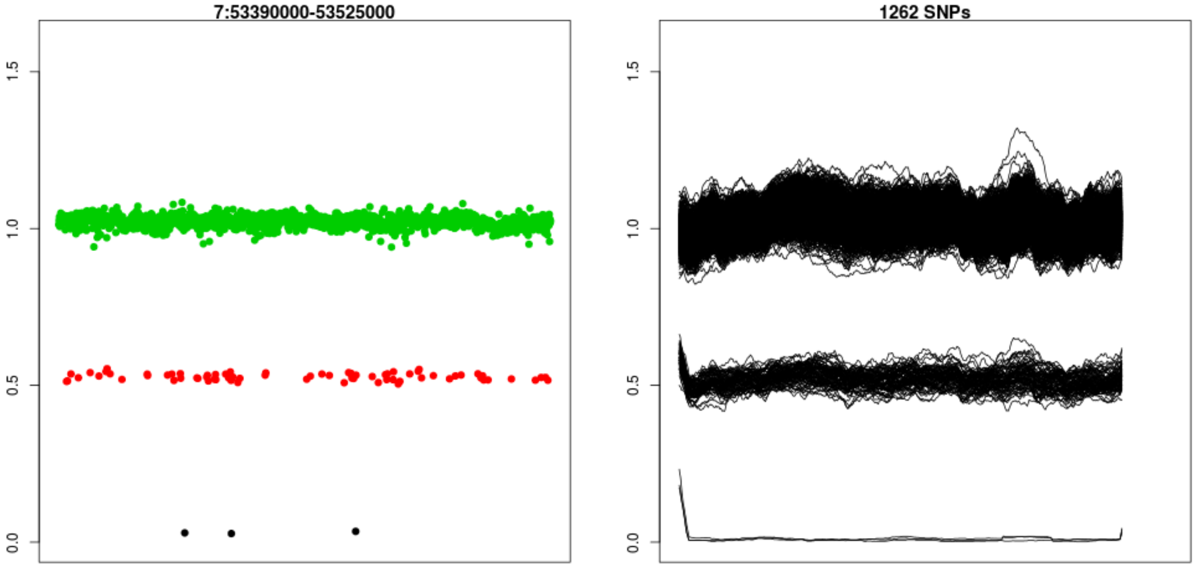

**Supplementary Figure 6:** Overlap between the four cohorts after segment-based and genotype quality control.

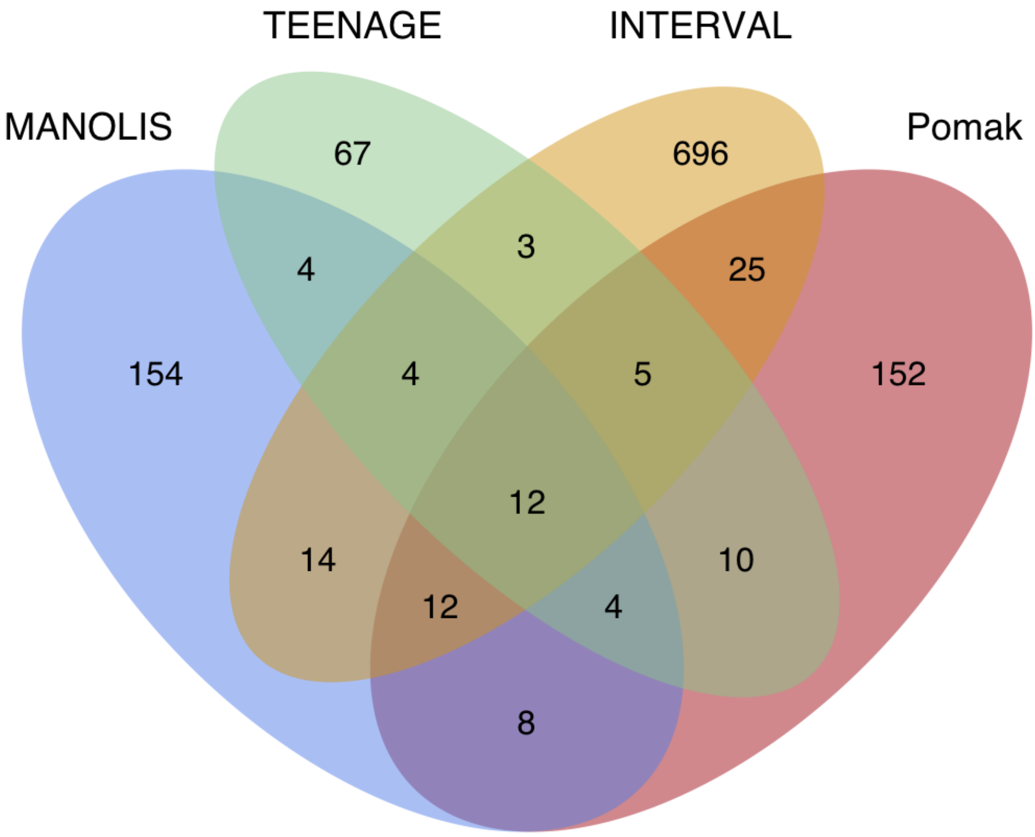


**Supplementary Figure 7 :** *CCL3* region and read depth. The dashed vertical line is the left boundary of the CNV called by UN-CNVc, which extends to the right of the displayed window. The shaded blue region is the CNV which was manually genotyped and gave rise to the association. Top plot: Red lines are the regressed depth segments belonging to individuals carrying a deletion of *CCL3L3*. The deletion is not localised to *CCL3L3* but extends far downstream, across a SNP-poor region. The duplication of *CCL3L3* by contrast is a much smaller event, which we used for determining the boundaries of the region to genotype. Protein coding genes and assembly exceptions are annotated below the plot. Second plot from top: Rolling average depth across 100 randomly selected samples confirms the complexity of this region. The *CCL4-TBC1D3B* duplication, the *CCL3L3* duplication and the *CCL3L3-TBC1D3F* deletion seem to be distinct events with differing depth patterns. The SNP-poor region downstream of *CCL3L3* also has lower average depth and SNP density, indicating a potentially difficult to access segment or a reference issue (third from top plot). This paucity of variants is reflected on the single-point association plot (bottom plot).The top associated variant is in strong LD with known *CCL3* pQTLs, and tags the deletion. (Figure on next page)

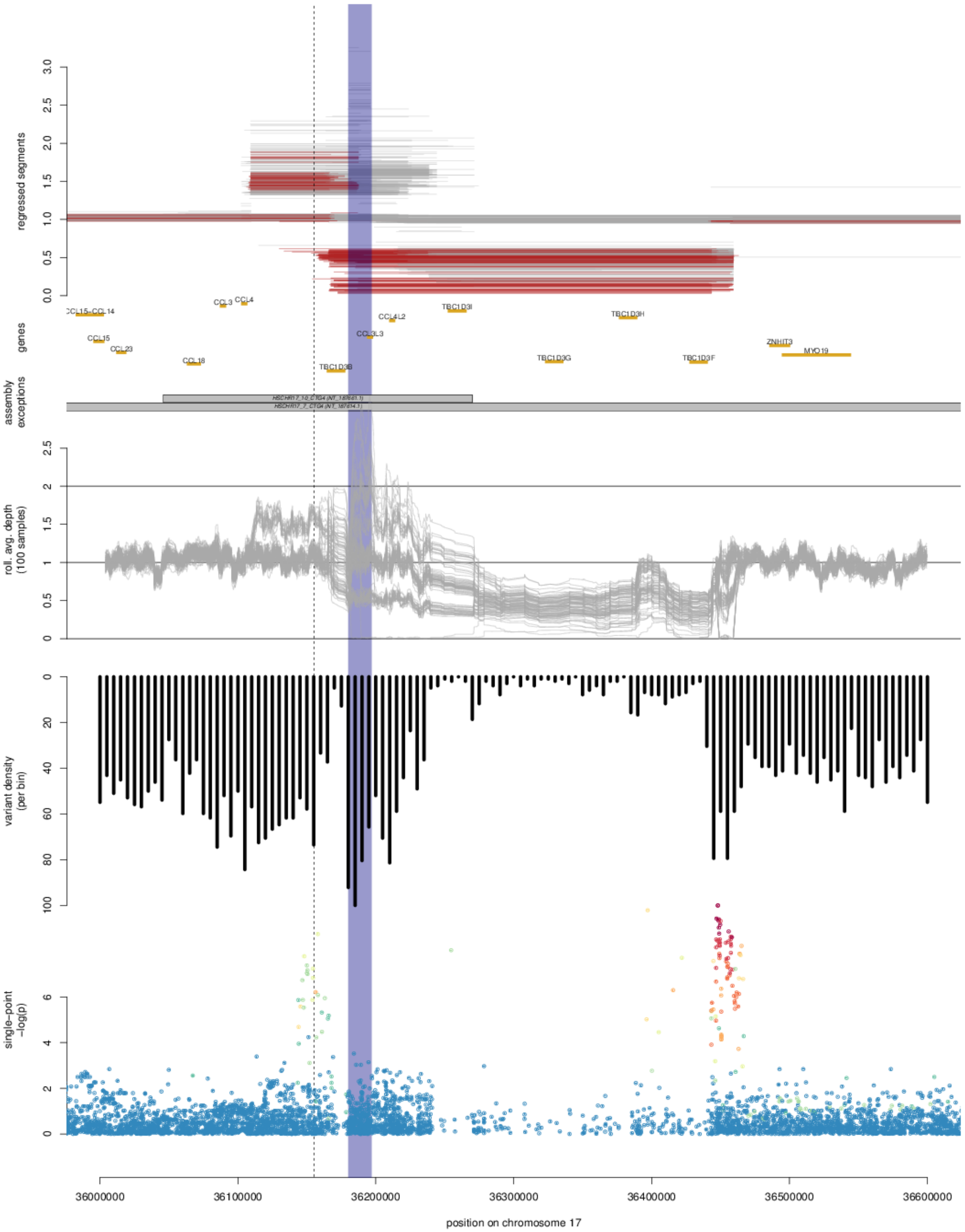

regressed segments
genes
CCL15-CCL14
CCL3
CCL4
CCL15
CCL23
CCL18
TBC1D3B
CCL3L3
CCL4L2
TBC1D3I
TBC1D3G
TBC1D3H
TBC1D3F
ZNHIT3
MYO19
assembly exceptions
roll. avg. depth (100 samples)
variant density (per bin)
single-point −log(p)
36000000
36100000
36200000
36300000
36400000
36500000
36600000
position on chromosome 17

**Supplementary Figure 8:** Association of *CCL3L3* deletion with CCL3 protein levels under the deletion and duplication (top) and deletion only (bottom) models. ANOVA are applied to called copy numbers, linear models are applied to uncalled relative depth.

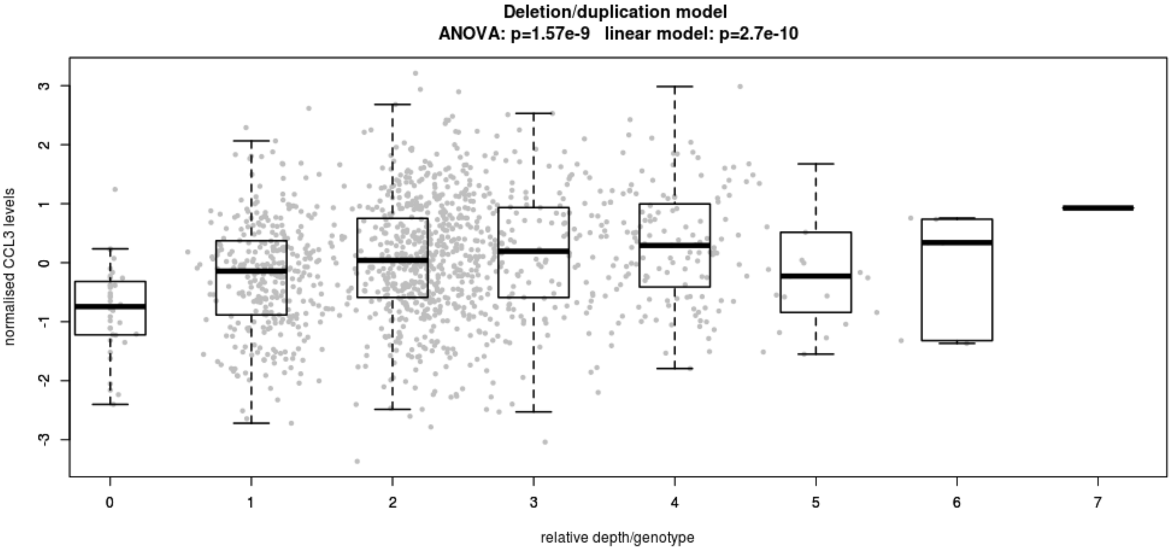


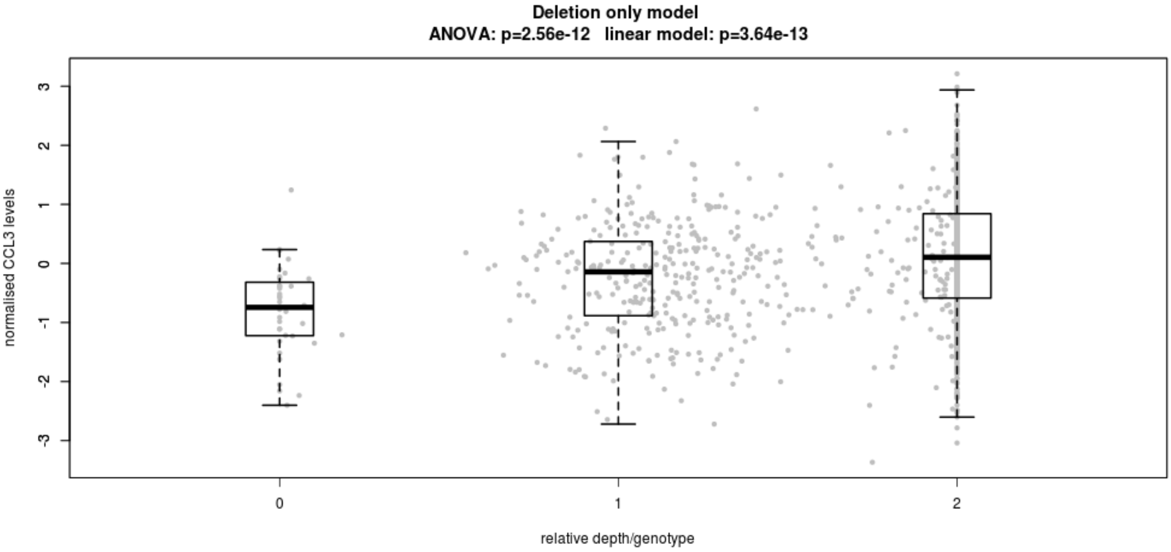

**Supplementary Figure 9:** Base 10 logarithm of the sizes of the CNVs called by UN-CNVc, GenomeStrip, and PennCNV.

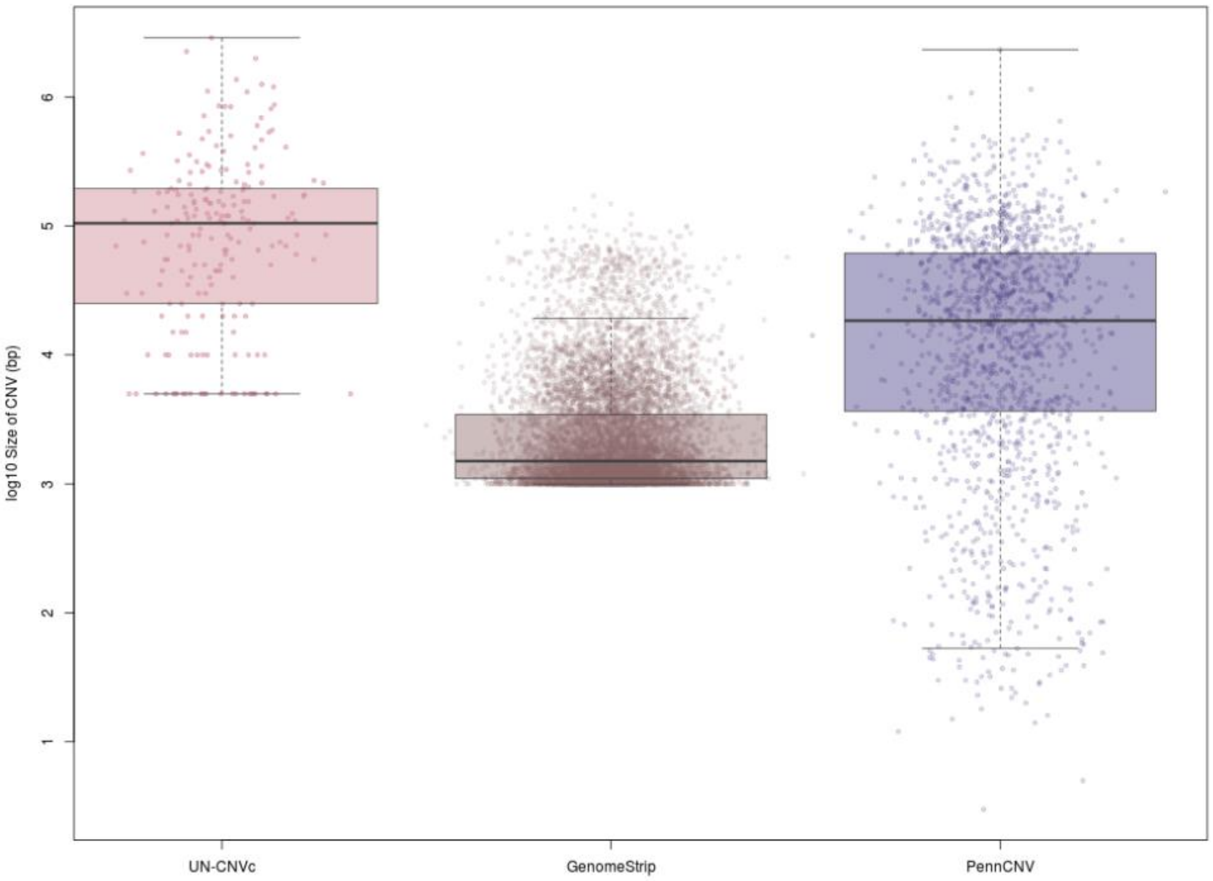

**Supplementary Figure 10: Influence of the complexity parameter *c* on depth regression.** Tested values of *c* are $1x10^{-4}$ for **a.**, $3x10^{-4}$ for **b.**, and  5x10-4 for **c.** The default used by UN-CNVc is the default of 0.01, with an option allowing the user to specify more sensitive values. Depth is measured on chromosome 11 in the Pomak cohort.

**a.**

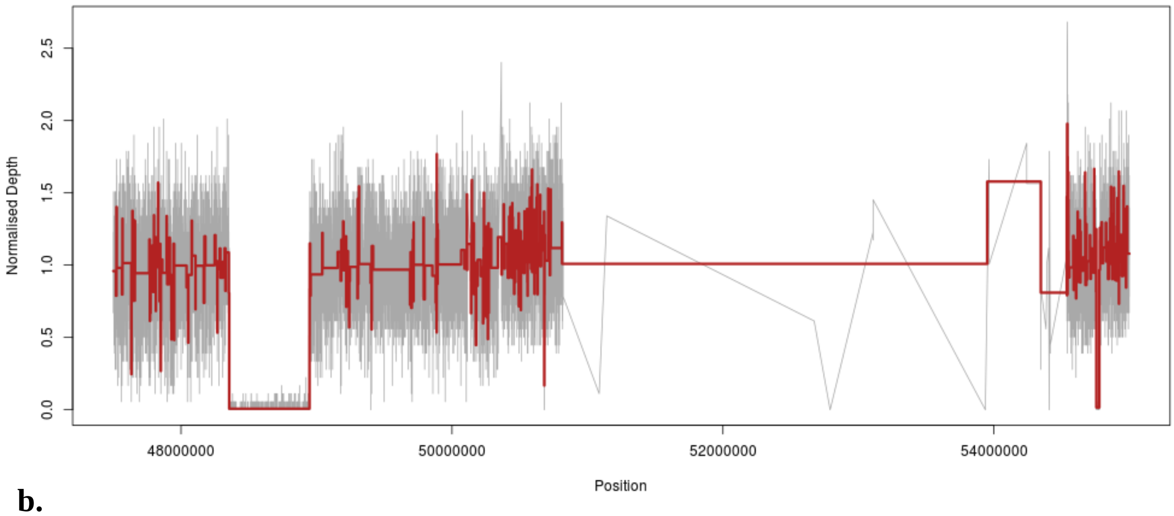


**b.**

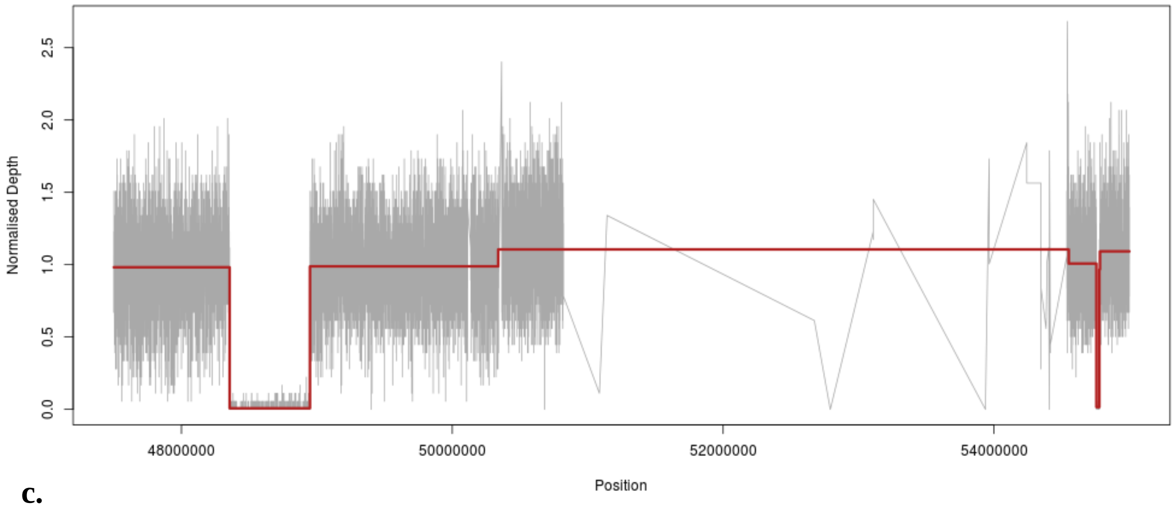


**c.**

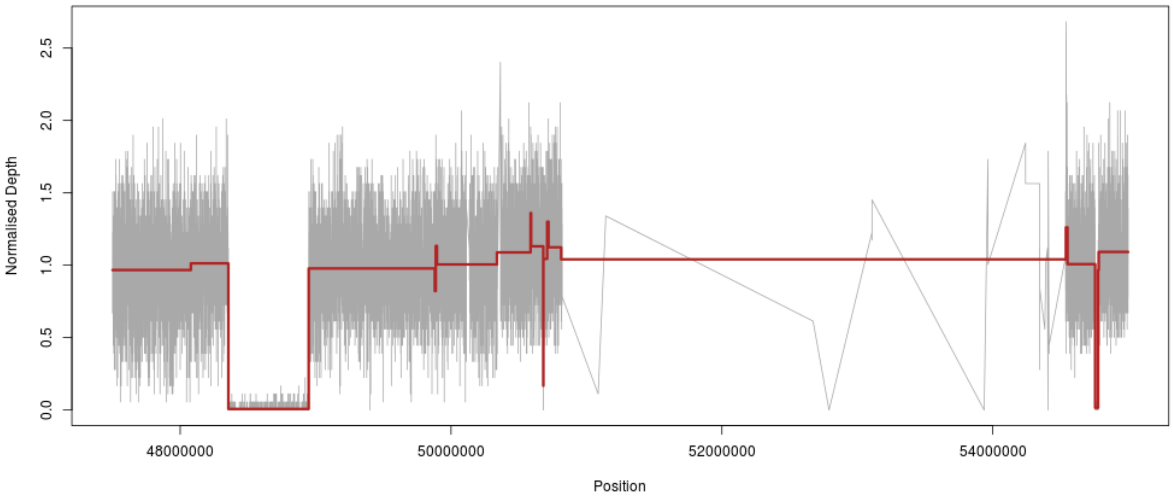
